# Context-dependent neural preparation for information relevance vs. probability

**DOI:** 10.1101/2024.05.20.594985

**Authors:** José M.G. Peñalver, Carlos González-García, Ana F. Palenciano, David López-García, María Ruz

**Affiliations:** Mind, Brain and Behavior Research Center (CIMCYC), University of Granada, Granada 18071, Spain; Data Science & Computational Intelligence Institute, Universidad de Granada, CP 18071

## Abstract

Preparation is a top-down phenomenon known to improve performance across different situations. In light of recent electrophysiological findings that suggest that anticipatory neural preactivations linked to preparation are context-specific and do not generalize across domains, in the current study we used fMRI to investigate the brain regions involved in these differential patterns. We applied multivariate decoding to data obtained in a paradigm where, in different blocks, cues provided information about the relevance or probability of incoming target stimuli. Results showed that the anticipated stimulus category was pre-activated in both conditions, mostly in different brain regions within the ventral visual cortex and with differential overlap with actual target perception. Crucially, there was scarce cross-classification across attention and expectation contexts except on a patch of the fusiform gyrus, indicating mostly differential neural coding of anticipated contents in relevance and probability scenarios. Finally, a model-based fMRI-EEG fusion showed that these regions differentially code for specific conditions during preparation, as well as specifically preparing for category anticipation in a ramping-up manner. Overall, our results stress the specificity of anticipatory neural processing depending on its informative role while highlighting a key hub of commonality in the fusiform gyrus.

## 1. Introduction

In recent times, cognitive neuroscience has experienced a resurgence of interest in proactive cognition, highlighting the central role of endogenous top-down brain processes. Within this framework, preparation can be conceptualized as an endogenous neural function that takes place prior to planned action and that improves subsequent behavior (Battistoni et al., 2017; González-García et al., 2016). It has been shown in several contexts, with marked interest in selective attention, or the ability to prioritize information relevant for behavior (e.g. Nobre & Serences, 2018) and expectation, or the generation of probabilistic predictions based on previous experiences (e.g. Schröger et al., 2015).

A large part of the investigation on attention and expectation has focused on their consequences on target stimulus processing (reviewed in Summerfield & Egner, 2009, 2016), which leads to contrasting effects on brain activation levels. While studies in human and non-human primates have found increased firing rates in neurons tuned to attended features (Chelazzi et al., 1998; Cohen & Maunsell, 2011; Kastner et al., 1999; Peelen & Kastner, 2011; Serences et al., 2004), studies of expectation classically show activation decreases (also known as expectation suppression, e.g. Kok et al., 2012a,b; Walsh & McGovern, 2018; see Feuerriegel et al., 2021 for a detailed review). Overall, results suggest that relevance and probability play different roles during target processing (Auksztulewicz & Friston, 2016; Gordon et al., 2019; Simon et al., 2018; Wyart et al., 2012).

However, at the anticipatory level, separate studies of attention and expectation have shown similar preactivation profiles across domains. Selective attention cues (Battistoni et al., 2017; Nobre & Serences, 2018) preactivate specific shape patterns in visual cortex (Stokes et al., 2009), relevant regions of spatial processing (Giesbrecht et al., 2006), as well as object- (Peelen & Kastner, 2011; Soon et al., 2013) and category-selective (Esterman & Yantis, 2010; González-García et al., 2018) perceptual regions. Relatedly, probabilistic cues lead to the preactivation of specific perceptual templates of oriented gabors (Kok et al., 2017), direction (Ekman et al., 2017), motor patterns (de Lange et al., 2013) or abstract shapes (Hindy et al., 2016). These findings have been extended and further detailed regarding specific stimulus categories such as human faces, which have been shown to be coded in face-selective regions during anticipation, with similar patterns as those induced by face perception, an effect that increases as the probability of perceiving a target stimulus increases (Blank et al., 2023). Importantly, when attention and expectation have been studied together it has usually been to define how both interact. For instance, it has been shown that during anticipatory stages, the representation of relevant stimulus categories is enhanced by probabilistic knowledge about certain characteristics of the target (Gayet & Peelen, 2022). However, the role of specific brain regions and how the underlying preactivated patterns differ across contexts of relevance and probability is currently unknown.

Altogether, although both attention and expectation seem to involve the preactivation of information, the effects on target processing are apparently opposed. This poses the question of whether preparation is a unified phenomenon or, conversely, reflects different mechanisms. Since previous evidence indicates that preparatory brain activity is related to stimulus processing (González-García et al., 2017; Hebart et al., 2018), it could be logically argued that different effects on target processing would be anticipated by different representational profiles that efficiently prepare the brain for relevant or probable target categories. However, making this claim asks for a direct comparison of the anticipatory preactivations in attention and expectation contexts. On this regard, a recent electroencephalography (EEG) study applying multivariate analyses successfully contrasted selective attention and expectations, crucially, during the preparatory interval (Peñalver et al., 2023). Anticipatory coding of incoming categories was found in both contexts, but the underlying neural codes did not generalize across them. The authors interpreted this by claiming that anticipatory templates are different across attention and expectation contexts. Results suggested that whereas anticipatory coding during attention was partially reinstated during target processing, coding appeared to be more abstract in the expectation condition. However, electrophysiological results are agnostic regarding the brain regions supporting these processes. Differences between relevance and probability anticipation could be due to separable anticipatory neural codes in the same regions, or activity in different brain areas. For example, anticipatory decoding could rely on object-selective regions in attention and lower order, less category-specific regions in expectation contexts.

Here, we used functional Magnetic Resonance Imaging (fMRI) to study the specificity of anticipatory representations in two key cognitive domains: relevance vs. probability. We adapted the task used in Peñalver et al. (2023) to fMRI, aiming at replicating and extending the findings that indicate that attention and expectation elicit different profiles of preparation. To do so, we performed a series of multivariate analyses tailored to study the representational characteristics of preparation and to disentangle different patterns of activity that may be taking place in similar regions. We then leveraged a model-based EEG-fMRI fusion approach to detail the temporal profile of preparation in crucial brain areas.

Based on the differences between attention and expectation observed during anticipatory (Peñalver et al., 2023) and target processing (e.g. Gordon et al., 2019; Zuanazzi & Noppeney, 2019), our overall hypothesis was that relevance and probabilistic preparation would lead to context-specific preactivations, which would translate into specific and non-generalizable anticipatory brain patterns in the attention and expectation conditions.

## 2. Methods

Methods are reported in accordance with the COBIDAS protocol (Nichols et al., 2016). The code used to preprocess and analyze the data is available at https://github.com/CIMCYC/MRI.

### 2.1. Participants

Forty-six participants (mean age = 21.98, range = 18-30; 23 women, 23 men) from the University of Granada were recruited and received 20-25 euros as compensation, depending on their performance. They were all native Spanish speakers, right-handed and with normal or corrected vision. The study was approved by the Ethics Committee for Research with Humans from the University of Granada, and all participants signed informed consent prior to participation. Besides, to comply with COVID-19 safety guidelines, they wore a face mask during the whole session, including the behavioral practice outside the scanner. Six additional participants completed the task but were discarded, two due to poor behavioral performance (<80% in any of the main conditions, attention or expectation), two due to excessive head movement (either over the voxel size or over 0.1° of translation and rotation in 2 runs or more) and other two due to technical issues during data collection. Sample size was calculated in advance to achieve a statistical power of 80% for an estimated small effect size (Cohen’s d = 0.3) and three within-subject independent variables (Block x Category x Cueing), and to match the one used in a previous experiment with a similar paradigm (Peñalver et al., 2023). Using PANGEA we estimated a minimum of 32 participants to detect the Block x Cueing interaction in reaction times and behavioral accuracy, our main behavioral prediction. Our final sample size (46 participants) provided an estimated power of 94% under the described parameters. Due to an incomplete orthogonalization of the cue-shape pairing in cases of movement in only one run, 2 participants were left out of some specific decoding analyses (n = 44).

### 2.2. Apparatus, stimuli, and procedure

Stimulus presentation and behavioral data collection were done with The Psychophysics Toolbox 3 (Brainard, 1997) on MATLAB (r2020) in a Microsoft PC. Stimuli were presented on an LCD screen (Benq, 1920x1080 resolution, 60 Hz refresh rate) over a grey background. The task, stimuli and parameters followed those used in our previous study (Peñalver et al., 2023) except that to adapt the task to fMRI we employed longer and jittered inter-event intervals, which reduced the total trial count. We employed 160 male and female faces (50% each, extracted from The Chicago Face Database (Ma et al., 2015) plus 160 unique Spanish male and female names (50% each). Four different geometrical shapes (circle, square, teardrop and diamond with thin black outlines, unfilled) were used as cues in the main task.

The task involved a paradigm in which cues provided information about the relevance (attention) or probability (expectation) of upcoming face or word targets (Figure 1). Half of the blocks belonged to the attention condition and the other half to the expectation condition. To control for perceptual confounds, two cue shapes were associated with faces and two with names (counterbalanced across participants). Note that these four cues were identical in attention and expectation blocks, and the nature of the cues (attention vs. expectation manipulation) was established by instructions appearing at the beginning of each block. That is, cues predicting faces in attention were the same as the ones predicting faces in expectation, and the same for names. Hence, within the attention condition participants saw four different cues (two for faces and two for names) combined in the different runs; the same combinations with same cues appeared in expectation. For each participant, cue pairs changed across the different runs although their predicted category (face or name) remained; the first cue for faces (e.g. a circle) appeared in half of the blocks with the first cue for names (e.g. a square) and the other half with the second cue for names (e.g. diamond). The task of participants was to indicate the sex/gender of the stimulus, responding whether or not the target belonged to the gender stated at the beginning of each block. Each block started with a screen in which they were informed about the block (attention or expectation), the target sex/gender (“Is the target male/female?”), and the two cues (one for faces and one for names). Importantly, the sex/gender was explicitly indicated only at the beginning of the block. The cues gave information exclusively about the incoming category of the target, either faces or names. Here we use the term “category” to refer to faces and names, regardless of gender. Given that attention and expectation are involved in almost any act of visual perception, we aimed at manipulating one process while keeping the other constant. In attention blocks, the cue indicated the relevant stimulus category to select (faces or names). Only if the stimulus belonged to the relevant category (50% trials, cued; e.g. is the target a female *face*?), the participant had to perform the gender discrimination task on the target. Otherwise, participants had to answer ‘no’ regardless of the stimulus sex/gender (non-relevant category, uncued). This manipulation of relevance, where further processing has to be applied only to selected stimuli, is similar to that employed in previous literature (e.g. Baldauf & Desimone, 2014; Saenz et al., 2002; Summerfield et al., 2006; Womelsdorf et al., 2006). Both relevant and non-relevant targets were matched in expectation, as by design they appeared with a 50% probability after each attention cue. In expectation blocks, the cue indicated the probable category of the target with a 75% likelihood (e.g. de Lange et al., 2013; Kok et al., 2017 for similar manipulations). Here, participants had to perform the gender discrimination task for both categories of stimuli, whether or not the target was cued (e.g. is the target a female *stimulus*?). This way, both the expected and unexpected targets were equally relevant. Participants were verbally instructed to use the cues in the two blocks to respond as fast as possible while avoiding errors. Importantly, the number of targets of each category in expectation was also 50% in all blocks, although they could appear with larger probability after being cued.

**Figure 1.**
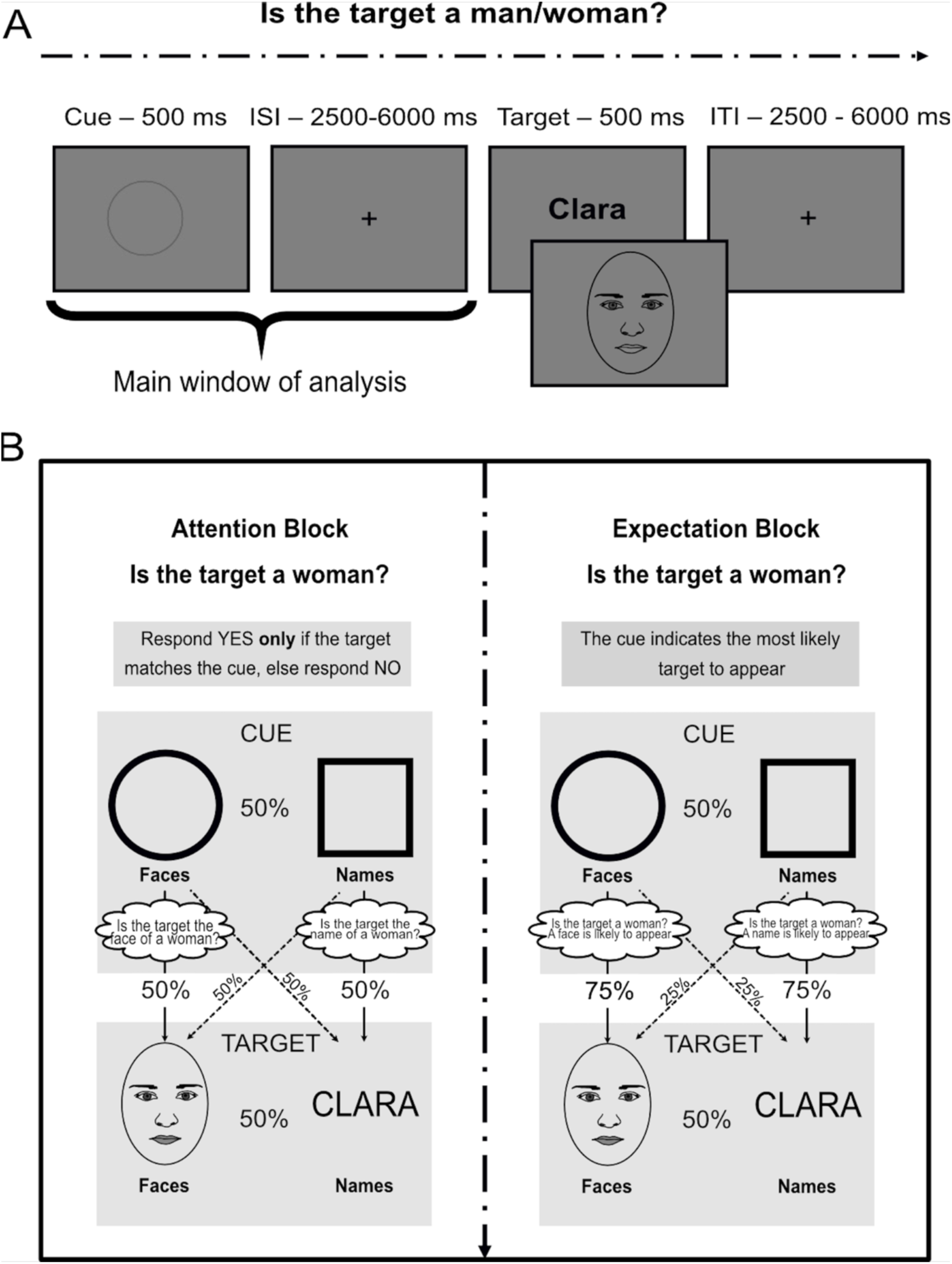
Behavioral paradigm. We employed a sex/gender judgment task embedded in a cue-target paradigm. Participants had to respond with an index finger to indicate whether or not the target belonged to the gender indicated at the beginning of the block. A) Schematic representation of a single example trial including presentation timings. B) Block design and summary of the possible stimuli. In each block there where two possible cues, which appeared with 50% probability in both attention and expectation runs. The cues were non-predictive in attention (where followed by each type of stimuli with a 50% probability), but correctly predicted the incoming stimulus category in expectation with 75% probability. Crucially, the number of targets of each category was also kept at 50% in both conditions. Classifiers decoded the predicted category in the cue interval or the actual stimulus in the target window. Note that although there were two cues per block, there were 4 shapes in total that changed throughout the blocks. Crucially, cues were identical across attention and expectation blocks.

In every trial of the main task, the sequence of events was as follows: a 500 ms cue (∼1.5°) was followed by a jittered CTI and then the target appeared for 500 ms (∼7°). Both the CTI and the inter-trial duration were jittered between 2500-6000 ms intervals, spaced in 700 ms steps following a uniform distribution (average 4250 ms). Each trial lasted on average 9.5 seconds and each run 7.6 minutes. The experiment was composed by 8 blocks (4 for attention and 4 for expectation) of 48 trials each, or 384 trials in total. Attention and expectation blocks appeared in separate runs in a fully alternated order, and the condition of the first block was counterbalanced across participants. Cues and target stimuli were also fully counterbalanced across participants. In total, the whole experimental session lasted 60 minutes approximately, plus additional practice outside the scanner.

### 2.3. Data acquisition and preprocessing

A single session of imaging was carried out with a 3T Siemens Prisma MRI scanner, equipped with a 64-channel head coil. T1-weighted anatomical images were obtained using a rapid acquisition gradient echo (MPRAGE) sequence (TR = 2250 ms, TE = 4.18 ms, TI = 900 ms, flip angle=9°, voxel size = 1 × 1 × 1 mm). In addition, two field map images (phase and magnitude) were collected to correct for magnetic field inhomogeneities (TR = 520 ms, TE1 = 4.92 ms, TE2 = 7.38 ms, flip angle=60°, voxel size = 2.5 × 2.5 × 2.5 mm). Whole-brain functional images were acquired using an echo planar imaging (EPI) sequence (TR = 1730 ms, TE = 30 ms, image matrix = 84 × 84, FOV = 210 mm flip angle=66°, slice thickness = 2.5 mm, voxel size = 2.5 × 2.5 × 2.5 mm, distance factor=0%, 50 slices with an acceleration factor of 2 (Simultaneous Multi-Slice acquisition). Slices were oriented along the AC-PC line for each participant.

The experiment consisted of 8 runs, each corresponding to a block of the behavioral task. For each run, 275 volumes were acquired, with the first 4 discarded from all runs. Anatomical images were defaced to ensure anonymization. MATLAB R2020 was used for preprocessing, which involved converting the raw DICOM images from the scanner into NIfTI files with BIDS format (Gorgolewski et al., 2017). For functional images, preprocessing (with SPM12 v7771) involved the following: (1) realignment and unwarping to correct for movement artifacts (using the first scan as the reference slice) and magnetic field inhomogeneities (using previously estimated fieldmaps); (2) slice timing correction; (3) corregistration with T1 using a rigid-body transformation and normalized mutual information cost function with 4th degree B-spline interpolation; (4) registration to MNI space using forward deformation fields from segmentation with 4th degree B-spline interpolation and MNI 2mm template space; (5) smoothing using an 8mm FWHM kernel. Multivariate analyses were performed with the unsmoothed, individual participant’s functional data space. Resulting images were later re-registered to the MNI space, smoothed and masked before second-level analyses.

### 2.4. Analyses

#### Behavioral

The task design had three within-subject factors: Block type (attention vs. expectation), Cueing (cued vs. uncued) and Stimulus category (faces vs. names). We calculated three-way repeated measures ANOVA for behavioral accuracy and reaction times (RTs) employing JASP (Love et al., 2019). For each participant and condition, trials with longer or shorter RTs than the average ± 2 SDs were discarded (11.5% on average). Since our data of interest for post-hoc analyses (cued vs. uncued expectation trials) did not meet normality criteria to perform t-tests, on neither accuracy (Shapiro-Wilk test, W = 0.91, p= 0.003), nor reaction time (W =0.92 p =0.004), planned comparisons are reported with Wilcoxon signed-rank test.

#### fMRI General Linear Model (GLM)

A GLM was implemented to estimate activity changes across conditions, and to obtain the beta coefficients to be used in subsequent multivariate analyses. We included condition-wise cue and target regressors in the model. Cue regressors were modeled using their presentation time (500 ms) plus the subsequent jitter for each trial. They were divided by Block (attention and expectation), Category prediction (Faces and Names) and Cue shape (Shape 1 and Shape 2). Importantly, although there was a total of 8 different cue regressors, these were distributed across the different runs. That is, in one run/block, cues could only be of a particular condition (e.g. Expectation), and the cues predicting a certain category had the same shape during the run (e.g. Face-shape 1 and Name-shape 2). Hence, each run included only 2 cue regressors. Target events were modeled as the presentation time of the stimulus on screen (500 ms), and consisted on regressors for the conditions of Block (attention and expectation), Category (faces and names) and Cueing (cued vs. uncued). Again, although there were 8 different regressors, they belonged to particular runs, which resulted in runs having 4 target regressors each. The model also included movement information of each participant, obtained during realignment. The regressors were convolved using the canonical hemodynamic response function (HRF). We estimated two sets of GLMs, one for univariate (with smoothed data) and another one for multivariate (without normalization and smoothing).

#### Mass Univariate Tests

We first carried out a univariate, voxel-level test to address the different activations in attention and expectation during the preparatory interval. To do so, we locked the events to the onset of cues, and directly contrasted both conditions by performing a paired t-test, collapsing across category and cue regressors. We performed this contrast for each participant, limited to grey-matter voxels (using the Neuromorphometrics atlas included in SPM, excluding white matter voxels), and then obtained statistical values from a second-level analysis, where they were compared using a one sample *t-*test against zero. Significance was established by first selecting voxels that passed a threshold of *p<*0.001, and the cluster size to a number of voxels corresponding with *p<*0.05, FWE-corrected (Eklund et al., 2016).

#### Decoding Analysis

Decoding was performed with The Decoding Toolbox (v 3.999F), using the beta images from the cue events obtained in the GLM. Classification was performed with the betas obtained from each block, rather than trial-wise, as this method provides larger signal-to-noise ratios, associated with higher statistical power (Allefeld and Haynes, 2014; see e.g. Díaz-Gutiérrez et al. 2020, Cristophel et al. 2015 for similar approaches). Moreover, since all trials within each of the eight anticipatory possible combinations (condition x category x cueing) had identical temporal structures, this choice avoided effects of collinearity when estimating trial-wise betas, which in turn lead to less stable and noisier beta coefficients (Abdulrahman and Henson, 2016). In all cases, we employed a two-class decoding approach with searchlight (sphere of 4 voxels radii, 251 voxels). A Support Vector Machine (SVM), with a leave-one-run-out cross-validation, was trained with all but one run, and then tested on the remaining one. This was repeated with all runs, and the results were averaged. To ensure unbiased classifications, we report balanced accuracies, which account for over-representation of one category over the other. Group statistics applied a one-sample *t-*test against zero. To correct for multiple comparisons, we identified individual voxels that passed a threshold of *p* < 0.001, and then the minimum cluster size was set to the number of voxels corresponding to *p* < 0.05, FWE-corrected.

With this approach, we performed two analyses. First, we classified attention vs. expectation. Since there is evidence of the involvement of frontoparietal regions in both attention (Dodds et al., 2011; Greenberg et al., 2010) and expectation (González-García & He, 2021), we intended to examine whether information in these regions is distinguishable between both conditions even if they generate similar activation levels. We used data from all cues (identical in shape and numbers across blocks) and both categories (faces and names) to train the SVM classifier. Since the attention and expectation manipulations appeared in different runs, for this analysis we grouped them in pairs of continuous blocks, obtaining four pseudo-runs that included one of each condition, and were suitable to leave-one-run-out classification. Importantly, this analysis is informative regarding the regions where the overall process of anticipation takes place differently depending on the relevance vs. probability manipulation.

Next, we queried the regions that carried specific anticipated content by performing the classification of predicted categories (faces vs. names) separately for attention and expectation. Specifically, for each condition, we classified all the cues (two shapes) that predicted faces versus all the cues (two shapes) that predicted names. Note that in each block the number of observations of each cued category was identical, making all blocks balanced. As in the previous section, the analysis was locked on cue regressors, modeled to include the entire anticipation jitter. Note that we did not compare cued vs. uncued targets, but cues that predicted (relevant or probable, depending on the block) face vs. word stimuli within the same block. Similarly to univariate analyses, to compare decoding levels across attention and expectation, for each participant we subtracted the results of the two conditions and applied a second-level one sample t-test.

#### Cross-decoding Analysis

Cross-decoding was performed to assess the extent to which different conditions shared coding patterns (Kaplan et al. 2015). That is, we trained a classifier in a particular condition, and then tested it on a different one. Significant above-chance classification suggests the existence of similar patterns of brain activity in the two conditions. The crossvalidation approach was adapted to each analysis to avoid confounds, which are detailed in the following paragraphs. Searchlight and second-level analyses were applied identically as in the two-class decoding, described above.

Our primary goal was to estimate the degree to which patterns of brain activity are shared for preparation for relevant vs. probable contents. We reasoned that if anticipatory attention and expectation recruit differential patterns of brain activity, category cross-classification should not yield any significant results. We trained the classifier with cues predicting faces vs. names in one condition (e.g. attention), and then tested it on the other one (e.g. expectation). This analysis was performed without applying cross-validation to increase the number of observations included in the analysis. Note that this approach, although not identical to the original analysis, does not entail any statistical limitation, since the datasets used for training and testing are independent.

The second goal was to examine pattern similarity between preparation and actual target perception. Thus, we trained the classifier to discriminate between faces and names with cue information, and then tested it on target data. We applied a leave-one-run-out cross-validation approach, by training with three cue runs, and then testing on the target of the remaining one. We did this separately for the cues in attention and expectation. We chose this direction of cross-decoding (train in the cue, test in the target) to ensure that the results showed similarity based on anticipatory representations, since target perception elicits a much larger level of activation engaged by sole stimulus input (targets or cues) which was not of interest (see Cichy et al. 2012 for a similar approach). Again, we compared attention and expectation cross-decoding results by subtracting participant-wise results and performing a second level analysis.

Finally, we employed a cross-decoding approach as a control analysis for perceptual confounds for the anticipation-based category analyses (Figure 4). Since there were two cues predicting faces and two predicting names, all of them in the two conditions, we trained the classifier in one pair (e.g. circle predicting faces and square predicting names) and tested it in the other (e.g. diamond predicting faces and drop predicting names), which should return information that is uniquely based in the content of the anticipatory categorical representations (see Peñalver et al. 2023 for a similar approach). We did this in the two directions and then averaged the results.

#### Regions of Interest (ROI) extraction

We formulated 5 ROIs in total, corresponding to the regions returning significant decoding results during the previous analyses. First, to study the consequences of better anticipated stimulus discriminability, we focused on the regions resulting from category decoding separately for attention and expectation during the preparatory window. Second, to study the source of the similarity between cue and target regressors, we employed the regions resulting from the cue-target cross-decoding analysis. Since this returned two distant enough clusters, one in the ventral visual cortex and another in the Frontal Operculum (FO), we split this result into two ROIs. Hence, the 5 ROIs were (1) bilateral Inferior Temporal Cortex (ITG), (2) occipital cortex, (3) left FO, (4) left anterior ITG and (5) occipital + temporo-ventral cortex.

ROI parcellation proceeded as follows. To avoid double dipping (Kriegeskorte et al., 2009), we performed a leave-one-subject-out procedure (LOSO, Esterman et al., 2010). For each participant, we repeated the second-level analysis while leaving that participant out of the sample, so that the particular ROI was not based on his/her own data. Then, the resulting clusters went through the same statistical correction described above. Finally, we registered the resulting ROIs back to each participant’s native space using the inverse deformation fields obtained during segmentation.

#### Voxel selectivity ranking

We applied a voxel selectivity ranking analysis to study whether neural tuning to different stimuli (instead of multivariate activity patterns) generalized from anticipation to stimulus perception. That is, whether the neurons that are most responsive to certain condition during anticipation are similar to those responsive to the same condition during target processing (see González-García & He, 2021; Richter et al., 2018 for similar approaches).

This analysis was performed on the 5 ROIs mentioned above. First, for each block type (attention or expectation) we obtained 8 different conditions to rank per each voxel. These were obtained from crossing category (faces and names) with runs (4 runs per block type). Then, for each ROI, participant and voxel, we obtained the betas associated with each condition as a measure of voxel activity during the cue, and ordered them from least to most activity induced. Next, for each voxel, we applied the same order obtained during anticipation to target perception. We reasoned that if voxel selectivity during the cue generalizes to the target window, the order of the eight conditions for each voxel when applied to the target should maintain a positive slope. Note that the order of the eight conditions depends on each voxel, and is not relevant by itself, but rather a means to sort target betas and study the corresponding slope. Next, we averaged all voxels of the ROI, obtaining a vector of the eight ranked values, and evaluated the slope of this vector by fitting a linear regression to the ranked parameter estimates. We obtained a slope value per participant, ROI and condition. Finally, we determined whether the slope was positive (and therefore suggesting generalization from cue to target perception) by performing a right-tailed one-sample *t-*test against 0, and then used False Discovery Rate (FDR) to correct for multiple comparisons. Then, if there was a positive slope in at least one of the two conditions (attention or expectation) we compared them using a two-tailed paired *t-*test, again corrected for multiple comparisons using FDR (González-García & He, 2021; Richter et al., 2018).

#### Model-Based EEG-fMRI Fusion

To investigate the spatial and temporal features of the findings, we employed an RSA-based EEG-fMRI Fusion (Cichy & Oliva, 2020; López-García et al., 2022). EEG data was obtained from a previous experiment (Peñalver et al., 2023) in which 48 participants performed the same task. The only differences were in the timing of stimulus presentation, since cues and targets were onscreen for 50 ms and 100 ms, respectively, and trial intervals were not jittered.

In both datasets, we employed cross-validated Mahalanobis distance (also known as Linear Discriminant Contrast, or LDC, see Walther et al., 2016) to estimate the dissimilarity (Kriegeskorte et al. 2008) between all combinations of condition (attention vs. expectation), category (cues predicting names vs. faces) and cueing (two cue shapes per predicted stimulus), rendering 8-by-8 Representational Dissimilarity Matrices (RDMs). LDC was chosen because it provides a continuous measure, making it highly informative and avoiding ceiling effects. Moreover, since it includes a cross-validation loop, we ensured that results were not affected by block-wise dependencies in either dataset. We calculated LDC following Bueno & Cravo, (2021). For every ROI (or time point) and each pair of conditions, we calculated the mean of each voxel (or EEG channel) splitting the data into two folds, a training and testing set. The distance between the two conditions was multiplied by the pseudo-inverse covariance matrix between the residuals of the first and the second conditions in the training set, and the distance values were then averaged across the two folds.

To perform the fusion, since participants were different in each experiment, we employed the group average of all RDMs in both modalities, obtaining one EEG-RDM per time-point, and 5 fMRI-RDMs, one for each of the ROIs (adapting the procedures in Hebart et al., 2018). For each time-point and region, we obtained the lower triangle of the symmetrical matrix, excluding the diagonal, and vectorized it for further calculations. The fusion value was the result of a squared Pearson correlation between the EEG-RDM and the fMRI-RDM, since this R^2^ coefficient is mathematically equivalent to the coefficient of determination of fMRI that explains the variance of EEG (Hebart et al., 2018). For each region, the procedure was repeated in every time point, rendering a time-resolved vector of R^2^ values for each ROI.

Although EEG-fMRI fusion reflects the spatial and temporal profile of the similarities between EEG and fMRI, it does not provide information regarding the cognitive information (in our case, condition, category or cue differences) that the results reflect. To obtain this, we employed a model-based approach (Hebart et al., 2018, Cichy & Oliva, 2020). First, we built three 8 by 8 theoretical RDMs, each one for a model, stating the predicted distances among conditions (see Supplementary Figure 1). These matrices were composed of 1s and 0s, with 1s reflecting large distances and 0s reflecting more similarity. The condition model predicts large distances between attention and expectation. The category model predicts large distances between cues predicting different categories, regardless of block or cue. Finally, the cue model predicts that conditions would be different when the physical shape of the cue differed. Then, we applied a commonality analysis (Seibold & McPhee, 1979) to study the variance shared by the different variables (in our case, EEG, fMRI and model RDMs, again following Hebart et al., 2018). We compared the two squared semi-partial correlation coefficients. One coefficient represents the shared variance between EEG and fMRI after controlling for all model variables except one variable (the model of interest) in fMRI. The other coefficient represents the shared variance when controlling for all model variables, including the model of interest, in fMRI. The variance shared between EEG and fMRI, explained by the variable of interest, is extracted by calculating the difference between these coefficients of determination (R^2^). We repeated this for every model, time-point and ROI. Importantly, to increase signal-to-noise ratio we smoothed EEG information by applying a 5-time point moving average. For every time point, the value was the average of that point, the two previous and the two posterior ones (see Peñalver et al. 2023 for a similar approach). Importantly, although correlation values can be either positive or negative, R^2^ is always positive, describing the overall relationship, but not the direction. However, commonality results respond to correlation, making it possible to find some negative values and results higher than the theoretical limit that is R^2^ (see Hebart and Baker, 2018 and Flounders et al. 2019 for similar cases). To make the interpretation of the results easier, in Figure 8 we represent the absolute value of the commonality analyses. Supplementary Figure 2 shows the original result.

In a first approach, we employed all the three models described above. Commonality (C) was calculated as follows:

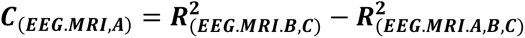

where A reflects the first model, B the second and C the third one. However, the block model explained a large proportion of the fusion results, possibly masking effects that could be accounted for by the other two models. Since regions were chosen because they revealed significant category decoding, we reasoned that condition (attention or expectation) differences could explain enough variance to mask any other interesting effects. Therefore, we repeated the analysis only with category and cue RDMs, to obtain a clearer representation of the possible underlying effects (see Flounders et al. 2019 for a similar approach). Note that data were still composed of two conditions, and hence we kept 8 by 8 RDMs. Two-model commonality is described as:

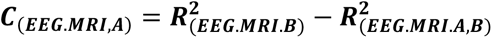

Statistical significance was calculated using a cluster-based permutation analysis. For each EEG-RDM, we permuted the data 5000 different times, obtaining as many permuted RDMs. Then, we repeated the analyses and obtained an empirical distribution of commonality values and cluster sizes. A window was considered significant if all the values lied over the 95% largest permuted results, and these were continued in more data points than the 95% of the cluster sizes. For similar approaches see Hebart et al. (2018), Flounders et al. (2019) and Peñalver et al. (2023).

## 3. Results

### 3.1. Behavioral

Participants’ overall performance showed high accuracies (M = 0.93, SD = 0.05). To assess behavioral effects, we used a three-way repeated measures ANOVA (Figure 2) on accuracy (Supplementary Table 1) and reaction times (Supplementary Table 2).

**Figure 2.**
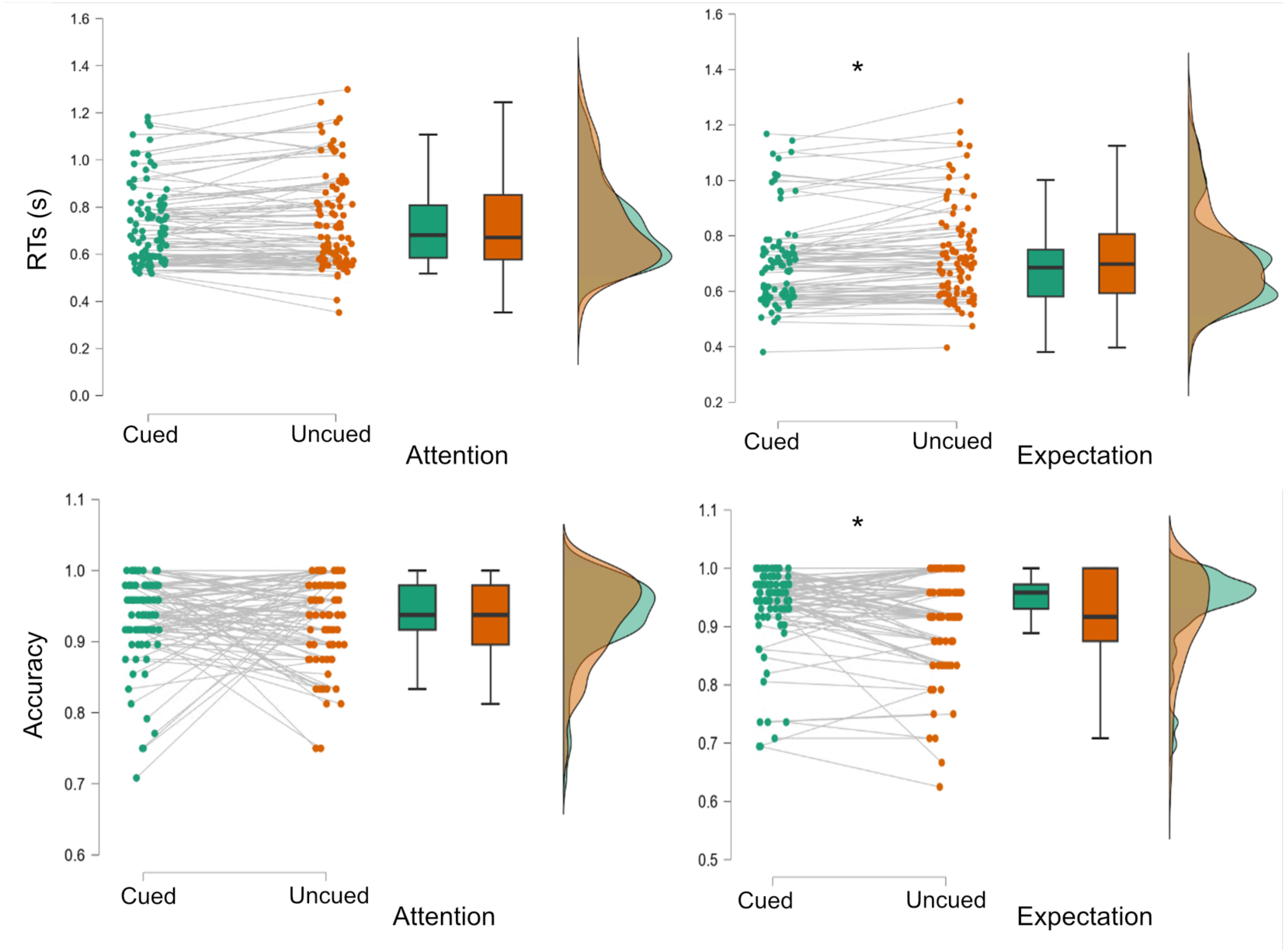
Behavioral results. Reaction times (top row, in seconds) and accuracy (bottom row) in attention and expectation blocks, for cued (green) and uncued (orange) trials. Dots represent individual participant’s scores per experimental condition. Grey lines connect each participant’s score in the two conditions of each block. The horizontal black line inside boxes represents the median, while the limits of the box indicate the first and third quartiles. Whiskers mark the 1.5 inter quartile range for the upper and lower quartiles. Lateral plots show the kernel distribution of each data value.

Behavioral accuracy only showed a main effect of Category (F_45,1_ = 13.02, *p* < 0.001, 𝜂p2 = 0.22), with less accurate responses to faces than to names (M = 0.92, SD = 0.06 vs. M= 0.94, SD = 0.07). Crucially, there was no main effect of Condition (F_45,1_ =0.31), indicating that attention and expectation did not differ in difficulty. Although the interaction between Block and Cueing was not significant (F_45,1_ = 1.98, *p=*0.16, see Supplementary Table 1 for the complete result), given our hypothesis of better performance in cued that uncued trials in the expectation condition, we performed the planned comparisons (using Wilcoxon signed-rank test) and observed better accuracies for expected vs. unexpected targets (Z_45,1_=2.53, *p* = 0.011, effect size = 0.43, M = 0.94 vs. 0.91), which was not found in the attention condition (Z_45,1_=0.58, *p* = 0.56).

Only the main effect of Cueing showed a significant main effect in reaction times (RT; F_45,1_ = 9.45, *p* = 0.004, 𝜂p2 = 0.17), as responses were generally faster in cued trials compared to uncued trials (M = 0.701 ms, SD = 0.17 vs. M = 0.732, SD =0.18). Again, although there was no Condition by Cueing interaction (F_45,1_ = 1.29, p=0.26), we studied the effect of expectations guided by our a priori hypotheses. Expectation trials showed faster responses for cued than for uncued trials (Z_45,1_=-4.41, *p* = 0.011, effect size = 0.7, M = 703 vs. 728 ms), which was not significant for attention (Z_45,1_=-1.58, *p* = 0.14).

### 3.2. Global differences between attention and expectation

Our first goal was to outline the regions that are involved in general anticipatory states during attention and expectation, either in global activations or in different patterns of activity. To do so, we first performed a univariate contrast for attention > expectation and vice versa, which revealed only small univariate differences during the preparatory window (Supplementary Table 3, Figure 3A).

**Figure 3.**
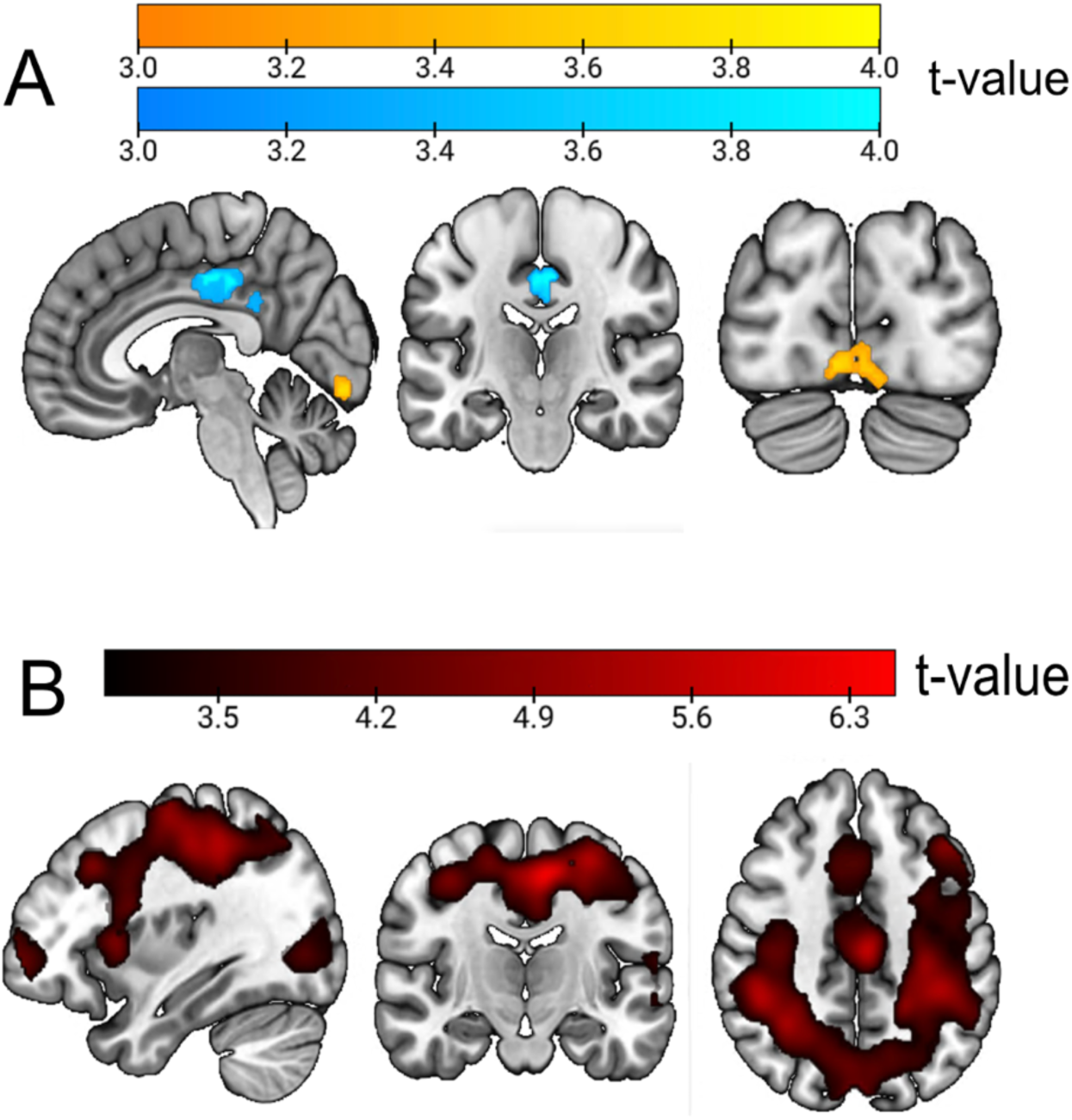
Global attention vs. expectation results during the anticipatory window. (A) Univariate GLM contrast comparing attention vs. expectation trials during anticipation. Scales reflect t-values. Yellow areas highlight significant clusters where attention induced larger activations than attention, while blue shows clusters where expectation induced larger activations than expectation. (B) Decoding results, classifying attention vs. expectation during the anticipatory interval. In both cases, minimum significant voxel threshold is t = 3.27.

Since previous literature has linked frontoparietal regions to anticipation in both conditions, we reasoned that even with similar univariate activation values, the patterns of voxel-wise activations should be different if attention and expectation lead to different effects. We studied this by performing a searchlight decoding of expectation and attention during the cue and preparation interval. Our results showed significant classification in two different clusters (Supplementary Table 4, Figure 3B). The first one included several frontoparietal regions associated with the multiple demand network (MD), such as the intraparietal sulcus (IS), Supramarginal gyrus (SMG), Superior Parietal Cortex (SPC), left anterior Insula and the Dorsolateral Prefrontal Cortex (DLPFC), with the peak voxel in the Supplementary Motor Area (SMA). The second one included broad visual regions, including the occipital gyri (OcG), with the peak voxel being at the left lingual gyrus (LiG). Note that both blocks were perceptually identical and were also equated in behavioral performance measures.

### 3.3. Differences in anticipatory category representations in attention and expectation

When looking at anticipatory activity separately for attention and expectation (Figure 4A [yellow – attention; blue – expectation] and Supplementary Table 4), we first observed significant coding of the anticipated relevant (attended) target category in two bilateral clusters in the Visual Ventral stream, with peaks at the left and right Inferior Temporal Gyri (ITG). In contrast, probable (expectation) category anticipations showed decoding mostly restricted to the early ventral visual cortex and Fusiform gyrus, with a peak at the Lingual gyrus. The comparison of these two results, by subtracting the maps of attention and expectation and performing a one-sample t-test against zero revealed that, although each condition peaked in different regions, decoding levels were overall not significantly different. This could be due to overall accuracies being low (as expected during anticipation) or to both conditions recruiting broad regions of the occipital and ventral cortices, although peaking in different areas. As mentioned before, these results are not driven by perceptual confounds, given that decoding was performed including two different cues for each category, which were differentially paired with the two cues of the other condition across blocks.

**Figure 4.**
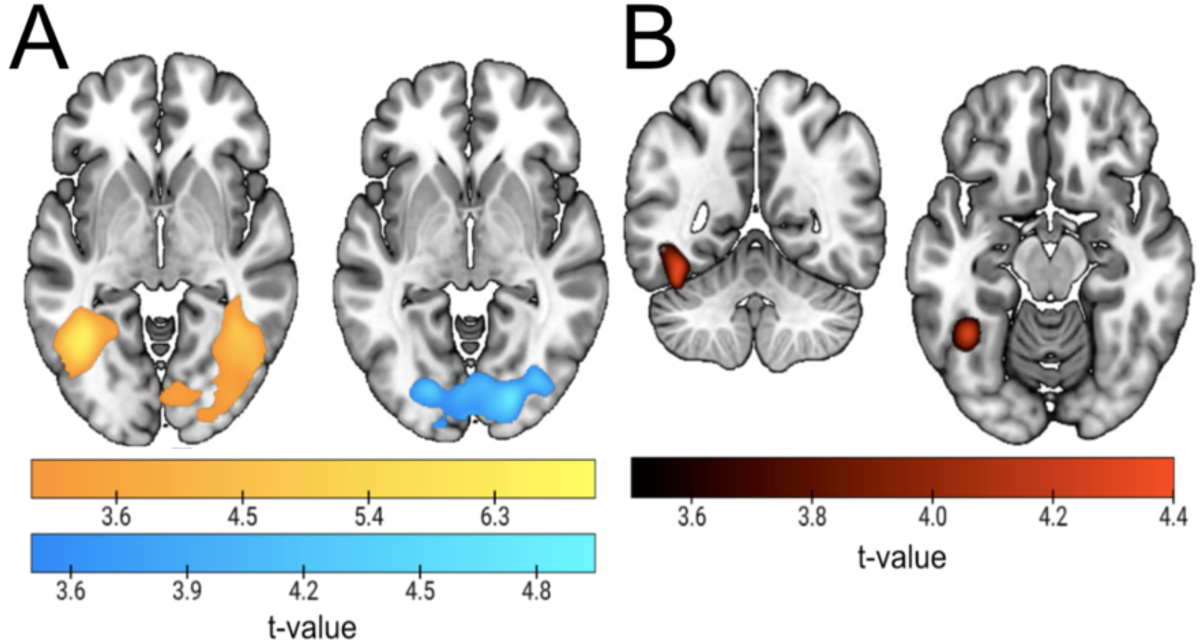
Classification results for anticipatory face vs. name decoding in attention and expectation blocks and cross-decoding among them. (A) Classification results in attention (yellow) and expectation (blue) blocks. Plot shows the result of classifying predicted/attended faces vs. names during the anticipatory interval. (B) Cross-classification results between attention and expectation. Figure shows the only significant cluster after training the classifier in one condition and testing in the other (and vice-versa, both directions averaged) during preparation. Note that the scales are different as they reflect different contrasts.

### 3.4. Category anticipation shows restricted generalization between attention and expectation

The previous results raise the possibility that preparatory neural codes are to certain extent shared between contexts of attention and expectation. To test for this, we performed a cross-classification analysis between relevance vs. probability anticipation. This analysis revealed a single cluster in the left FG (k = 756, Figure 4B and Supplementary Table 5).

### 3.5. Regions involved in anticipation partially overlap with those related to target perception

Results in the previous two sections show that category anticipation in attention and expectation engages similar but not completely overlapping regions, which employ mostly different coding patterns across contexts. This could be due to anticipatory representations in both conditions differing in their level of similarity with target perception. Therefore, we investigated the potential pattern overlap between preparation and the actual perception of face and name stimuli, separately for attention and expectation. To do so, we trained the classifier during cue processing and tested it during target processing (Figure 5 and Supplementary Table 5; the result of averaging of the two training-testing directions for the two conditions can be found in Supplementary Figure 1). In attention blocks (Fig. 5A, yellow), the left anterior ITG and left Frontal Operculum (FO) showed common patterns for both processing stages. For expectation (Fig. 5A, blue), results were limited to the bilateral Occipital Gyri (OcG), peaking at the left Inferior Occipital Gyrus (IOG). A conjunction analysis only returned a cluster in the left FG, matching the coordinates of the significant regions of the previous cross-classification analysis (Section 3.4; Figures 4B and 5B). We compared both conditions by performing a t-test, which returned an occipital bilateral cluster where cue-target cross-decoding was largest for expectation (Figure 5C and Supplementary Table 5). This suggest a larger involvement of early visual areas in the coding of probable stimuli that is specific to the cognitive context of expectation.

**Figure 5.**
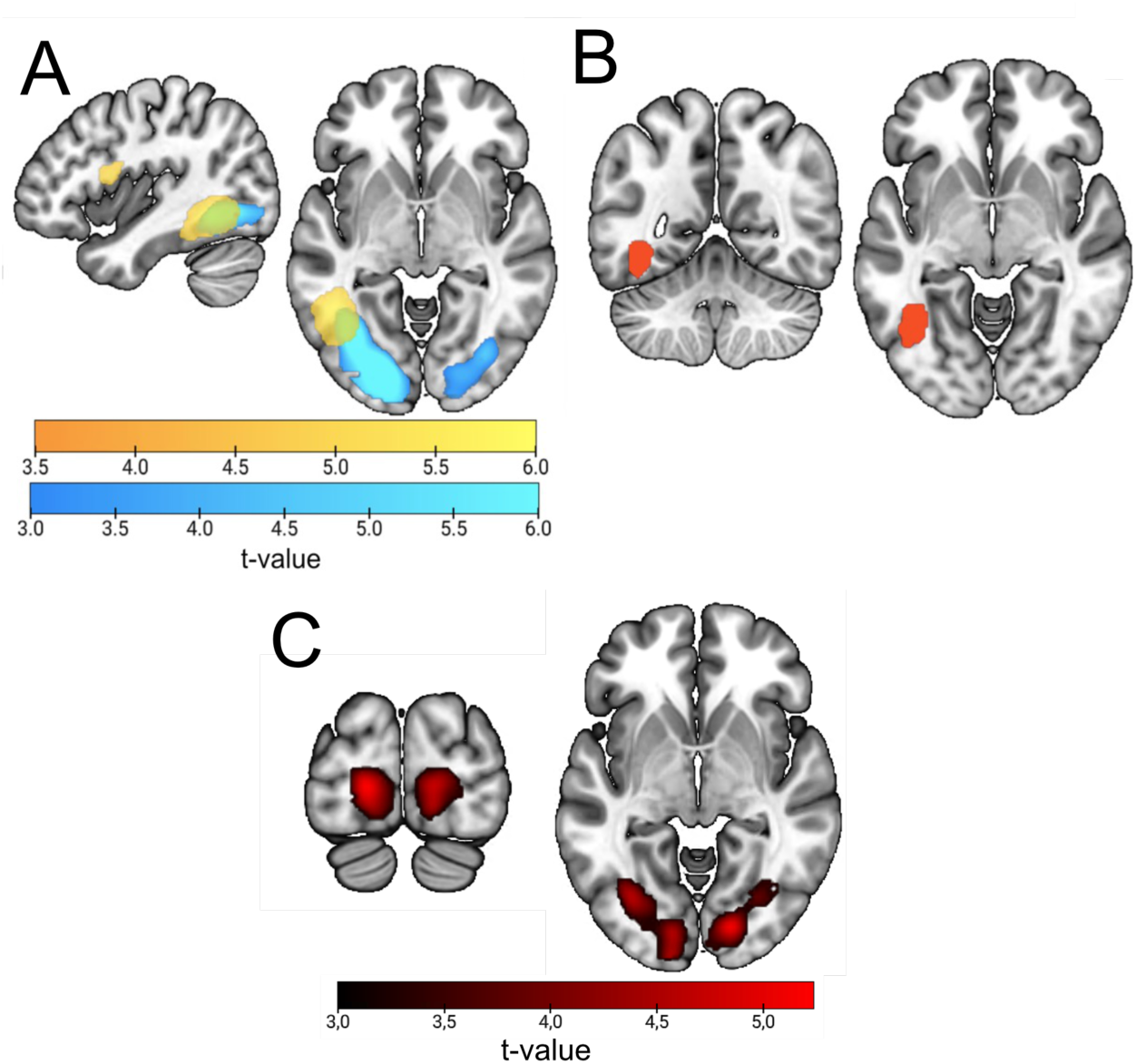
Cue-Target Cross-decoding results. (A) Results of training the classifier to differentiate faces vs. names during anticipation and testing it during stimulus perception (target). Attention results are shown in yellow, and expectation in blue. (B) Voxels that are significant in the two conditions in the analysis in A (binarized conjunction). This matches the overlap between the attention and expectation clusters in Figure 5A. Moreover, note the remarkable similarity between this result and Figure 4B. (C) Contrast between cross-decoding results in attention vs. expectation. Depicted are the areas where the expected stimulus category is significantly better cross-decoded than the attended one, after a second level analysis contrasting decoding accuracies of both conditions shown in (A). There were no significant clusters in the opposite direction.

A mechanistic explanation for the similarities found between preparatory and perceptual coding might be accounted for by neurons in both epochs of the trial being tuned to similar stimuli. We studied the stability of neural tunning from cue to target stimuli in a univariate manner, which also allowed direct comparisons between attention and expectation. We established a voxel selectivity ranking during anticipation, and regressed it onto image preference during the target. We performed this analysis in the five ROIs obtained from category decoding during the cue, and cue-target cross-decoding. We separately analyzed the slope of each condition (see Figure 6 for results). The slope was only significantly positive for expectation, in the ventral visual ROI associated with cue-target cross-decoding for expectation (t_43,1_=3.27, *p* < 0.001). Moreover, the result was also significantly more positive than in the attention condition (t_43,1_=2.86, *p* = 0.004). This result suggests that voxels within early occipital and temporo-ventral regions are tuned in a similar way during cue and target epochs, but in expectation trials, in line with the previous cross-decoding results. Note that while the result in this ROI could appear to be driven by larger differences between conditions in the most selective condition a control analysis using only the first seven points was also significant (attention slope = −0.005, expectation slope = 0.077, t = 2.5, p = 0.02).

**Figure 6.**
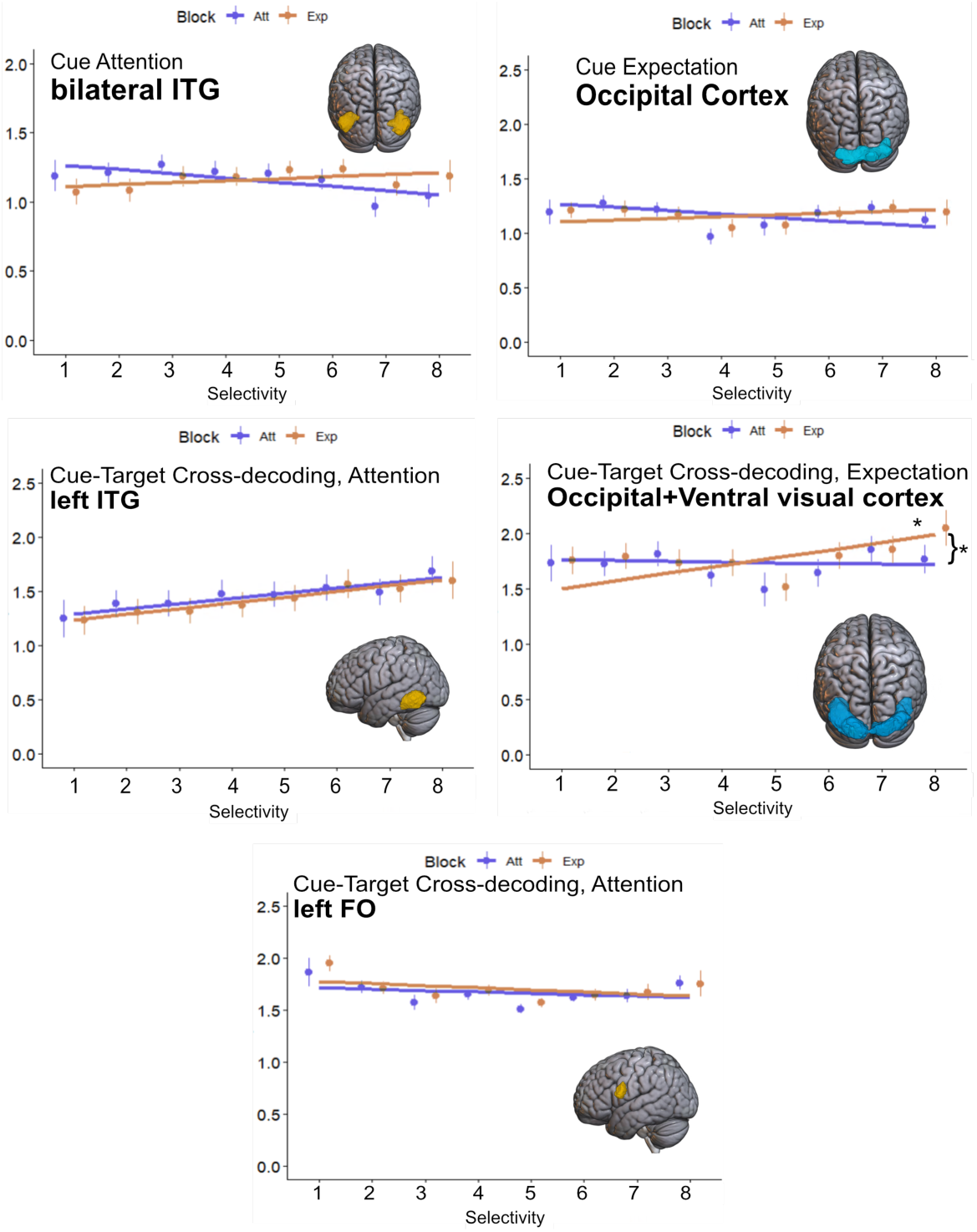
Voxel selectivity results. Each plot depicts the results for one ROI. The Y axes show average beta values across all voxels and participants. The X axes show selectivity preference from least (one) to most (eight) of the eight conditions during anticipation, which was then applied to target activity (see methods). Dots represent mean voxel and participant beta values, and vertical lines the SD. Continuous lines indicate the slope obtained after fitting a linear regression to each condition. Asterisks indicate statistical significance. Yellow ROIs are the ones obtained from attention blocks results, while blue ROIs come from expectation blocks. ITG = Inferior Temporal Gyrus; FO = Frontal Operculum.

### 3.6. EEG-fMRI fusion shows category and block coding in ventral regions

Model-based EEG-fMRI fusion results (Figure 7, grey shaded area) showed similarity between EEG and fMRI data that in most regions increased with time, with the exception of the most occipital area (ROI 2). This result stands out in the left FO (ROI 3), where similarity is large during the whole interval. Crucially, the most ventral regions showed evidence of similarity that was less present in occipital areas. This might suggest that this area represents anticipated contexts with high detail but with a relatively late latency.

**Figure 7.**
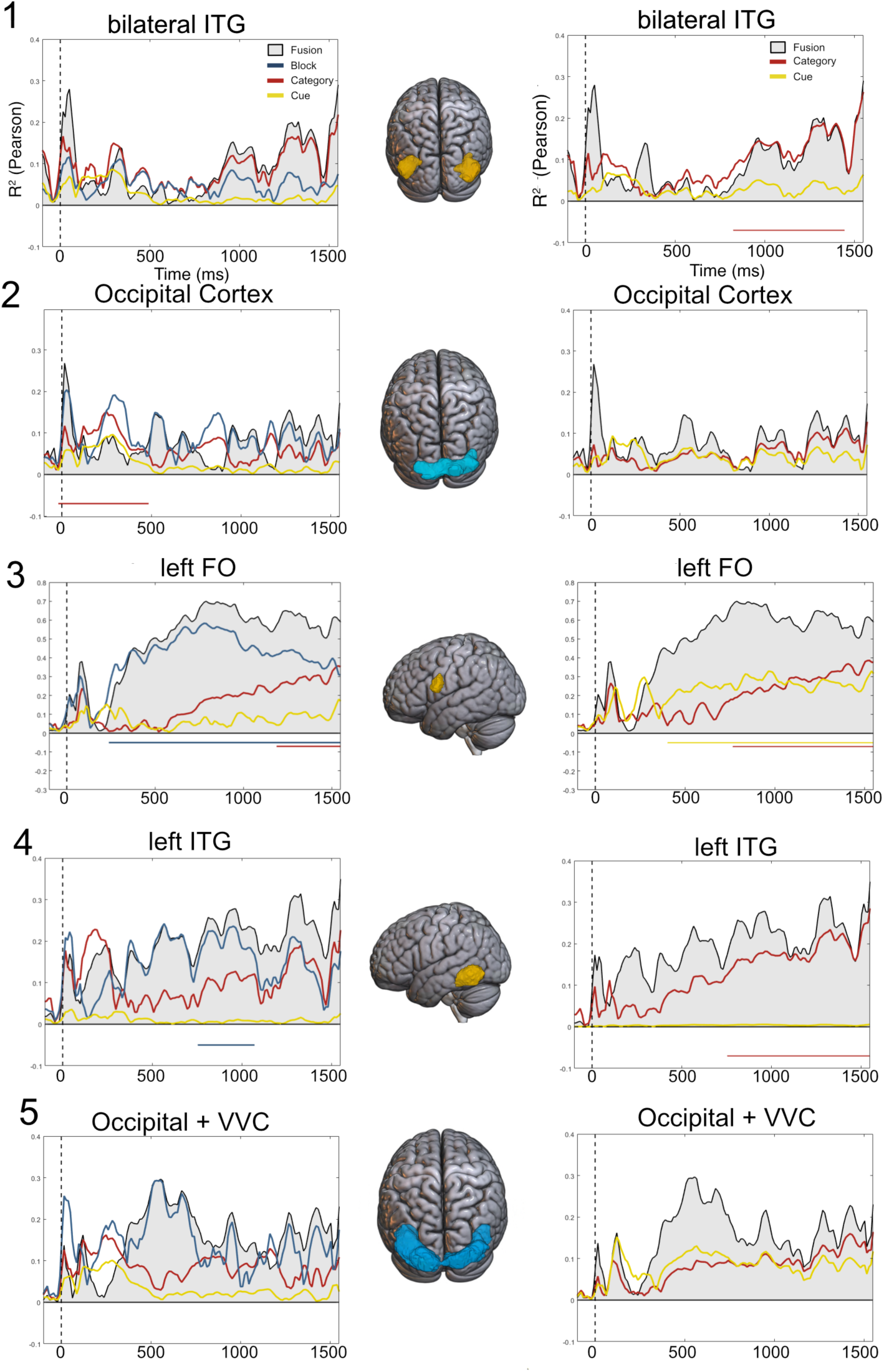
Model-based EEG-fMRI fusion results. Fusion (grey area and black outline) results. Y axes show R^2^ values, and X depicts time. Vertical dashed line indicates cue onset, while the 1550 ms limit is the target onset in the EEG experiment. The left column shows commonality results for three models: block (blue), category (red) and cue (green). The right column shows commonality for two models, category and cue. Significance appears in horizontal lines below the axis, colored as the corresponding significant model. Note that fusion values are identical in both columns. The two first ROIs are extracted from significant clusters of the decoding of face and name anticipated targets, yellow in attention and blue in expectation. The second pair comes from cue-target cross-decoding, following the same color pattern. Note that the scale of the Y axis on ROI 3 is larger. ITG = Inferior Temporal Gyrus; FO = Frontal Operculum.

To investigate the cognitive functions subserving the fusion results we applied commonality analyses (Figure 7 – left column). We designed three RDMs based on three models accounting for condition (attention vs. expectation), category (face vs. name) and cue (the actual shapes of the cues), depicted in Supplementary Figure 2. The analysis revealed that most of the similarity between EEG and fMRI patterns was explained by the condition model, especially in the FO from 200 ms to the end of the preparatory interval. This suggests a sustained pattern of differences in attention and expectation, that was common across both experiments and that is maintained while anticipating incoming information. However, these regions were selected because they showed a significant level of category decoding in previous analyses, which is also the main focus of this investigation. We reasoned that the variance explained by the condition model might have masked relevant information pertaining to the category model. Hence, we performed a second commonality analysis, this time only including category and cue models. Results are shown in Figure 7 – right column.

In this latter analysis the category model explained a significant part of the variance in three out of the five regions. Moreover, the pattern of commonality in all of them, except in the occipital cortex, described a ramping-up tendency, akin to the one found for similar analyses in Peñalver et al., 2023. Interestingly, the FO also showed evidence of cue shape representation, which was not present in more visual areas, suggesting the involvement of this region in the whole anticipatory process, including shape interpretation. Overall, ventral regions showed persistent coding of the contextual condition, which was largest in more anterior the region was. However, category anticipation was also present in more occipital areas. Overall, fusion analyses showed that block differences were sustained in visual regions, while the anticipated category was represented during most of the interval, peaking before target onset.

## 4. Discussion

The current study characterizes the specificity of the anticipation of relevant (selective attention) vs. probable (perceptual expectations) information content. Our findings show that top-down preparation is highly specific, evidenced by relevance and probability contexts leading to distinguishable brain states that extend from cue to target processing. Moreover, although anticipated stimulus categories are coded in the brain before their actual presentation in both contexts, the underlying preparatory patterns are mostly unique for attention and expectation. Overall, these results challenge the notion that preparatory templates in selective attention and perceptual expectation are equivalent, favoring a view instead where the informational role of mental contents shapes their underlying neural patterns.

Our first goal was to study whether attention and expectation contexts lead to different anticipatory states. We found small univariate differences in preparatory coding of different task sets (González-García et al., 2018; Hebart et al., 2018; Peñalver et al., 2023). Multivariate results extended these findings, showing that although activation levels were similar in attention and expectation in frontoparietal regions (Greenberg et al., 2010; Summerfield et al., 2006), their subserving patterns differed. Note that these results cannot be explained by differences in perceptual factors or task difficulty, since perceptual details of the cues and behavioral results in attention and expectation were equated. We found two clusters, one comprising a wide range of left occipital areas, extending from early visual regions to the left posterior ventral cortex, and another one in frontoparietal sites typically associated with the Multiple Demand network (MD), including the anterior insula, FO, SPC and MFC (Dosenbach et al., 2007; Wen et al., 2018). This network has been related to attention and cognitive control, memory load (Manoach et al., 1997) or task switching (Wager et al., 2004). Overall differences in our task are likely related to the contextual mechanisms recruited to anticipate demands and their relevance vs. probability informational function in the selection of the appropriate response (e.g. Woolgar et al., 2015).

Crucial to our research was to investigate whether the representational nature of relevant or probable information about incoming stimuli was also different. Results showed that it was possible to decode the anticipated stimulus category (faces vs. names) in both contexts, but the regions associated with this effect differed for the most part. The bilateral ITG represented anticipated information in relevance contexts. This region has been associated with object recognition, including faces (Verhoef et al., 2015) and words (Willems et al., 2016), also in anticipatory settings (Willems et al., 2016). The expectation manipulation, in contrast, led to category decoding in earlier visual regions including the lingual gyrus, associated with initial visual processing (Mechelli et al., 2000). Interestingly, attention and expectation differed in the anterior-to-posterior location of the anticipatory representations, with the former being in higher regions of the hierarchy, while the latter engaging earlier perceptual sites. This is similar to what was found by Kok et al. (2016), where a spatial attention manipulation had larger effects on higher order visual areas (V2, V3) while expectation was detected in V1. Also in line with these findings, a recent paper by Garlichs & Blank (2024) suggested that expectations influence face processing not only in face-selective regions, but also in the occipital face area, matching our expectation results. Specifically, they showed how the expected attributes of faces were sharpened in that region, while more ventral areas showed a dampened profile, interpreted as higher reliance on prediction errors than on priors. Extrapolating these features to the anticipatory window in our experiment, it could be the case that probable information induces patterns of high fidelity with broad features of the incoming stimulus, while more ventral areas where information may be attenuated (e.g. to avoid redundancies) do not represent the information in such a clear manner, making it more difficult for the classifiers to pick up such information. In the same line, attention is thought to sharpen stimulus processing in the ventral stream (Goddard et al., 2022; Woolgar et al., 2015). It could be possible that areas sharpening relevant target features exhibit decodable patterns during the anticipatory interval. Noteworthy, our analyses classified faces vs. names, which canonically engage different areas (Kanwisher et al., 1997; Mechelli et al., 2000), similar to other studies that have employed faces and houses (Ragni et al., 2021). While it is possible that the decoding observed reflects the engagement of different regions, rather than patterns in the same areas across conditions, it is unlikely that this is the primary driver of our results, given that control univariate contrasts comparing anticipation for faces and names did not return significant differences in the anticipatory interval. Moreover, anticipatory decoding was found in broad visual regions, including V1, which would unlikely be differently activated by different stimulus categories. Further developments could employ more similar stimuli, such as different grating orientations (e.g. Kok et al., 2017) to establish whether differential anticipatory representations take place also for stimuli that are perceptually similar.

In addition, to query the similarity of neural patterns across conditions, we implemented a cross-classification analysis between attention and expectation blocks. There was evidence of cross-decoding, but it was limited to a cluster in the left Fusiform Gyrus. This could suggest that the anticipation of specific relevant vs. probable stimulus categories could be partially, although not completely, based on different mechanisms, replicating and extending previous results (Peñalver et al., 2023). Interestingly, the cluster in the left FG suggests that part of the coding during anticipation is shared across contexts, and provides further evidence in favor of preparation sharing characteristics in attention and expectation contexts, while, at the same time, being subserved by distinct patterns of brain activity.

A potential source for differences between attention and expectation could be that they engage different levels of overlap with actual target perception, similarly to what has been suggested for visual imagery (e.g. Cichy et al., 2012). Cross-classification between cues and targets was evident in both conditions, however these effects appeared in mostly different areas. Attention showed cross-decoding on the left ITG, plus in the IFG. On the other hand, this analysis revealed more anterior occipital and temporal ventral regions for expectation. Surprisingly, significant regions did not fully match the ones found during anticipated category decoding, which suggests that anticipatory coding was taking place also in these areas but did not reach statistical significance levels. Interestingly, a small region in the FG appeared in both attention and expectation (see Figure 5B), which almost exactly matched the cross-classification cluster shared in anticipation across contexts. Hence, the two analyses together bring forward the notion that at least part of the preparatory neural codes are shared across conditions, despite the specificity of the requirements. Altogether, this suggests the left FG might be a key hub to code relevant and probable incoming information similarly in otherwise different preparation contexts.

A voxel selectivity ranking analysis provided further evidence regarding the degree of generalization of neural patterns from cue to target stimuli. We did this in the regions related to category anticipation in previous analyses: cue decoding and cue-target cross-decoding for attention and expectation. Out of the five regions, we only found evidence of generalization in the cue-target region for expectation, in broad ventral stream sites. There, only expectation showed evidence of generalization, and it was also significantly larger than for attention. Hence, this set of results highlights that probabilistic preparation for different categories, compared to context of relevance, is more akin to stimulus perception.

As a last step, we detailed the temporal profile of the fMRI results through a RSA-based fusion (Cichy & Oliva, 2020; Flounders et al., 2019; Hebart et al., 2018) in the ROIs obtained in previous analysis. This approach is also relevant to provide further support to the true anticipatory nature of the findings obtained with the fMRI methodology. Although the paradigm jittered the time intervals between cue and targets to minimize overlap in the signal, the sluggish nature of the BOLD response hinders a clean separability of both task epochs. However, this is not an issue with the EEG data, which has excellent temporal resolution. Results of the fMRI-EEG fusion provide the temporal profile of the activation patterns in the selected ROIs. These indicated that while the ITG showed sustained similarity with EEG activity, more occipital regions were associated with specific peaks at the middle and the end of the preparatory window, which might indicate specific transient reinstatement of the anticipated information. A further model-based commonality analysis revealed that the majority of these results were accounted for by differences between attention and expectation. Furthermore, when studying only the category and cue models of anticipation, we found that the anticipated category explained the similarity between EEG and fMRI patterns during most of the preparation interval, and in a similar ramping-up fashion as the one found in Peñalver et al. (2023). Our results further reveal that this ramping-up pattern takes place in visual regions, but also engages regions of the Frontal Operculum, which could be acting as a source of bias of more perceptual regions (e.g. Baldauf & Desimone, 2014). Ramping-up mechanisms have been suggested to serve working memory in reinstating useful information in contexts of temporal anticipation (Barbosa et al., 2020; Jin et al., 2020; Ruz & Nobre, 2008), an explanation that fits with our findings. However, further research is needed to clarify the computational role of these patterns, and their differential role across contexts.

Our results add to the evidence that attention and expectation engage neural preparation, aligning with predictive processing accounts of the anticipatory effects in both contexts (Auksztulewicz & Friston, 2016; Feldman & Friston, 2010; Kok et al., 2017). Attention consistently showed anticipatory coding in regions of the ventral visual cortex that are related to categorical stimulus perception, while expectation was related to earlier visual areas. In this scenario, selection could be operating on representations of the relevant stimulus categories, perhaps more closely related to brain imagery (Christophel et al., 2015; Lawrence et al., 2019). This could happen by preparatory increases of gain in neurons tuned to relevant categories, which could enhance information sensory weights during target processing (Feldman & Friston, 2010). On the other hand, expectation could induce excitability increases in earlier regions tuned to more basic stimulus features, allowing for a more flexible representation and induction of prediction errors to reduce noise during visual processing. Importantly, this is further supported by the voxel selectivity ranking results showing how voxel tunning remained stable only in expectation, adding to the notion of neural excitability and responsiveness to probable stimuli. However, our results also show that attention and expectation do engage at least partially overlapping mechanisms, as shown by cue decoding and cue-target cross-decoding analyses. One brain region, located in the left fusiform cortex, showed overlapped coding of incoming stimulus categories in attention and expectation. This suggests that anticipation conforms a set of mechanisms, with distinctive while also shared representations in some key neural hubs.

It is also important to note that the manipulation of the relevance or probability induced by the cues in our paradigm necessarily led to differences in task settings that may have affected the results observed. It could be the case that anticipatory representations *per se* were similar across conditions, but the specific task requirements highlighted specific aspects of these patterns, leading to overall contextual differences picked by the classifiers. For example, in the expectation condition participants could neglect the cue and prepare for the overall gender category queried, whereas this was not possible in the relevance condition. However, two key results make this possibility unlikely. First, behavioral observations showed an effect of expectation, such that when the predicted category (faces or names) differed from actual target stimuli, responses were slower and less accurate. This could only happen if participants prepared for face and name categories. Moreover, cue and target intervals cross-generalized in object selective regions such as the fusiform gyrus when classifying both names and faces, suggesting that these two categories were differently represented in both conditions. Nevertheless, we cannot fully dismiss that contextual task requirements had an effect on our results; further studies should attempt at ensuring fully independent manipulations.

Another potential limitation is that perceptual cue details could have confounded the anticipated category decoding performed separately in the attention and expectation conditions, in the sense of decoding the shape of the cues instead of their meaning. This is however unlikely. First, we trained and tested the classifiers using the same set of four cues, which having a varied set of shapes reduces the likelihood of perceptual details driving the classification (see Etzel et al. 2016 for a similar approach). Additional control analyses training the classifier in a pair of cues and testing on the other (and thus fully removing perceptual differences) provides similar results, although only for attention, potentially due to a lack of statistical power (see Supplementary Figure 3 and Supplementary Table 5). Besides, cross-classification between attention and expectation was substantially limited and absent in lower-level perceptual regions, which speaks against perceptual decoding since cue shapes were equal between the two blocks. Also, there were different regions subserving attention and expectation anticipatory coding; had classifiers be using perceptual features, these regions should show larger levels of overlap. Additionally, classifiers trained on the cue generalized to the target epochs in broad higher order perceptual regions. Crucially, the visual identity of the cues was fully independent of the targets employed, and cues were far less perceptually salient than targets, making it unlikely that the result were solely due to the cue’s perceptual features. Moreover, the cue anticipation results obtained with the EEG data in our previous study, which cross-classified with fully independent cue shapes with excellent temporal resolution (Peñalver et al. 2023), provide reliable fusion indexes with the fMRI cue data, strengthening the idea of similar underlying anticipatory processes.

Finally, in the present study we performed decoding analyses using block-based betas instead of trial-wise outputs, thus reducing the number of observations for decoding analyses. While the subject has been debated (e.g. Arco et al. 2018), we chose to use block betas because they provide larger signal-to-noise ratios compared to single-trial estimates, which increases power in decoding analyses (Allefeld and Haynes, 2014) while significantly reducing taxing computational requirements, that scale with sample size. Moreover, since all trials within each of the eight anticipatory possible combinations (condition x category x cueing) had identical temporal structures, this choice prevented effects of collinearity when estimating trial-wise betas, which in turn lead to lead to noisier beta coeffecients (Abdulrahman and Henson, 2016). Note also that the lack of cross-classification was even more drastic in our previous EEG dataset, which included a much larger number of observations in all the classifications performed (Peñalver et al. 2023).

Altogether, with a paradigm that manipulated the anticipation of relevant or probable contents in tasks equated perceptually and in difficulty levels, our neuroimaging results show that the preactivation of specific patterns in anticipation of task demands is a complex phenomenon that entails mainly unique mechanisms in each context, also with some neural substrates shared across attention and expectation. In both conditions cues pre-activated patterns similar to those reinstated during subsequent target perception. However, this overlap happened in mostly different regions, and only in expectation it was possibly due to specific neural tuning to probable stimuli. Our findings suggest that selection acts through complex stimulus representations, while expectation increases excitability in earlier, more basic perceptual regions. Overall, our results replicate and extend previous findings, thus stressing the specificity of anticipatory processing depending on the informative role it plays for further target processing.

## Data and Code Availability

All the code necessary to preprocess and analyze the data used in this manuscript, and a description of how to produce each figure can be found at https://github.com/ChemaGP-UGR/AttExp_fMRI while EEG and fMRI data are at OpenNeuro: https://openneuro.org/datasets/ds005386/versions/1.0.0

## Author Contributions

José M.G. Peñalver: Conceptualization, Data curation, Formal analysis, Writing – original draft. Carlos González-García: Writing – review & editing, Supervision. Ana F. Palenciano: Formal analysis, Writing – Review. David López-García: Formal analysis, Writing – Review. María Ruz: Conceptualization, Resources, Writing – review & editing, Supervision, Project administration, Funding acquisition.

## Acknowledgements

This research was supported by grants PID2019-111187GB-100, PID2022-138940NB-I00 funded by MICIU/AEI/10.13039/501100011033 and FEDER, UE, awarded to MR. JMGP was supported by a post-doctoral contract granted by the University of Granada and funded by grant RYC2021-033536-I to CGG. CGG was supported by Project PID2020-116342GA-I00 funded by MCIN/AEI/10.13039/501100011033, and Grant RYC2021-033536-I funded by MCIN/AEI/10.13039/501100011033 and by the European Union NextGeneration EU/PRTR. We are grateful to Marta Becerra Losada for her assistance with data collection.

