## Supplementary Materials for "Context-dependent neural preparation for information relevance vs. probability"

**Supplementary Table 4.1.** Behavioral accuracy results.

| Conditions | F | p | $\eta^2_p$ |
|---|---|---|---|
| Block | 0.311 | 0.580 | 0.007 |
| Cueing | 2.231 | 0.142 | 0.047 |
| Category | 13.024 | < .001 | 0.224 |
| Block ✻ Cueing | 1.977 | 0.167 | 0.042 |
| Block ✻ Category | 24.112 | < .001 | 0.349 |
| Cueing ✻ Category | 6.766 | 0.013 | 0.131 |
| Block ✻ Cueing ✻ Category | 6.717 | 0.013 | 0.130 |

**Supplementary Table 4.2.** Reaction Time results.

| Conditions | F | p | $\eta^2_p$ |
|---|---|---|---|
| Block | 1.475 | 0.231 | 0.032 |
| Cueing | 9.451 | 0.004 | 0.174 |
| Category | 1.679 | 0.202 | 0.036 |
| Block ✻ Cueing | 1.291 | 0.262 | 0.028 |
| Block ✻ Category | 1.417 | 0.240 | 0.031 |
| Cueing ✻ Category | 0.679 | 0.414 | 0.015 |
| Block ✻ Cueing ✻ Category | 5.384 | 0.025 | 0.107 |

**Supplementary Table 3.** Univariate contrast results

| Contrast | Region | Coordinates | Cluster size (k) | Z | p | T(peak) |
|---|---|---|---|---|---|---|
| Cue Att.> Cue Exp. | V2 | 8, -80, -12 | 216 | 3.62 | 0.04 | 3.94 |
| Cue Exp.> Cue Att. | rpCC | 2, -16, 40 | 367 | 3.9 | 0.004 | 4.3 |
| | lpCC | -4, -42, -28 | 210 | 3.84 | 0.044 | 4.22 |

Notes: Region labeling based on the Julich-Brain atlas (Amunts, 2020). P-values are cluster values corrected for multiple comparisons. V2 = Secondary visual area, rpCC = right posterior Cingulate Cortex, lpCC = left posterior Cingulate Cortex. Att. = attention, Exp. = expectation.

**Supplementary Table 4.** Decoding accuracy results.

| Contrast | Region | Coordinates | Cluster size (k) | Z | p | Decoding (peak) |
|---|---|---|---|---|---|---|
| Cue Att. vs. Exp. | SMA | -6, -20, 50 | 27841 | 5.46 | <0.001 | 60.34% |
| | LiG | -6, -76, -6 | 2302 | 3.86 | <0.001 | 57,69% |
| Target Att. vs. Exp. | PRG | 34, -18, 52 | 54668 | 6.40 | <0.001 | 62.21% |
| | STC | -48, -40, 2 | 1595 | 4.93 | 0.002 | 56,44% |
| Cue category decoding – Att. | lITG | -42, -58, -6 | 1564 | 5.73 | 0.001 | 59.91% |
| | rITG | 42, -58, -10 | 1439 | 4.84 | 0.001 | 57.31% |
| Cue category decoding – Exp. | LiG | -20, -84, -8 | 2761 | 4.4 | <0.001 | 57.46% |

Note: Decoding accuracy (balanced accuracy) shows the accuracy value in the peak voxel. SMA = Supplementary motor area, LiG = lingual gyrus; PRG = Precentral Gyrus, STC = Superior Temporal Cortex, ITG = inferior temporal cortex, l and r = left and right.

**Supplementary Table 5.** Cross-decoding results

| Contrast | Region | Coordinates | Cluster size (k) | Z | p | Decoding (peak) |
|---|---|---|---|---|---|---|
| Cue category between blocks | lFG | -42, -50, -22 | 756 | 4.01 | 0.026 | 54.13% |
| Cue-Target category – Train Cue - Att. | lITG | -48, -52, -8 | 1246 | 5.29 | 0.002 | 57.13% |
| | lFO | -52, 6, 20 | 594 | 4.17 | 0.03 | 54.17% |
| Cue-Target category – Train Cue- Exp. | lIOG | -38, -62, -2 | 8235 | 6.64 | <0.001 | 57.12% |
| Cue-Target category – Exp. – Att. | V1 | 18, -86, 0 | 2544 | 4.59 | <0.001 | T = 5.26 |
| Cue-Target category – Bidirectional - Att | lITG | -48, -52, 8 | 1291 | 5.3 | 0.002 | 57.23% |
| | lFO | -50, 4, 20 | 595 | 4.1 | 0.033 | 54.05% |
| Cue-Target category – Bidirectional - Exp | lIOG | -38, -62,2 | 8602 | 6.59 | <0.001 | 57.2% |
| Cue category across different cues - Att | rFG | 36, -60, 8 | 1248 | 4.93 | 0.002 | 55.36% |
| | S1 | -40, -22, 44 | 1199 | 4.07 | 0.002 | 56.3% |

IFG = Inferior Frontal Gyrus, lITG = left Inferior Temporal Gyrus, lFO = left Frontal Operculum, lIOG = left inferior occipital gyrus, lFG = left Fusiform Gyrus. rFG = right Fusiform Gyrus. Att. = attention, Exp. = expectation. S1 = Primary Somatosensory Cortex.

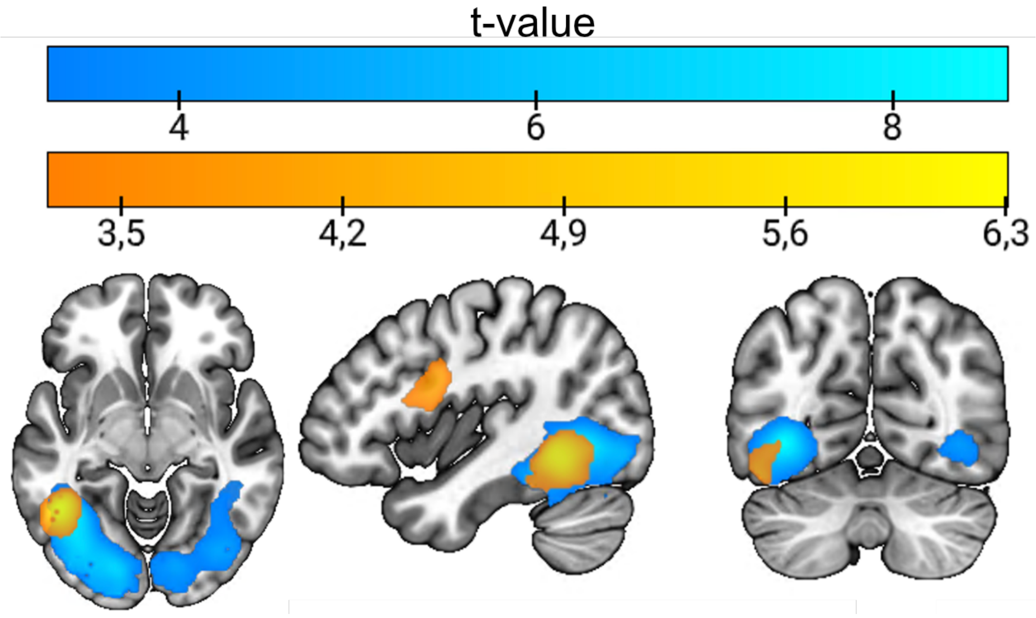


**Supplementary Figure 1.** Cue-Target Cross-decoding results, average of training and testing in the two two directions (cue-target and target-cue). Attention results are shown in yellow, and expectation in blue.

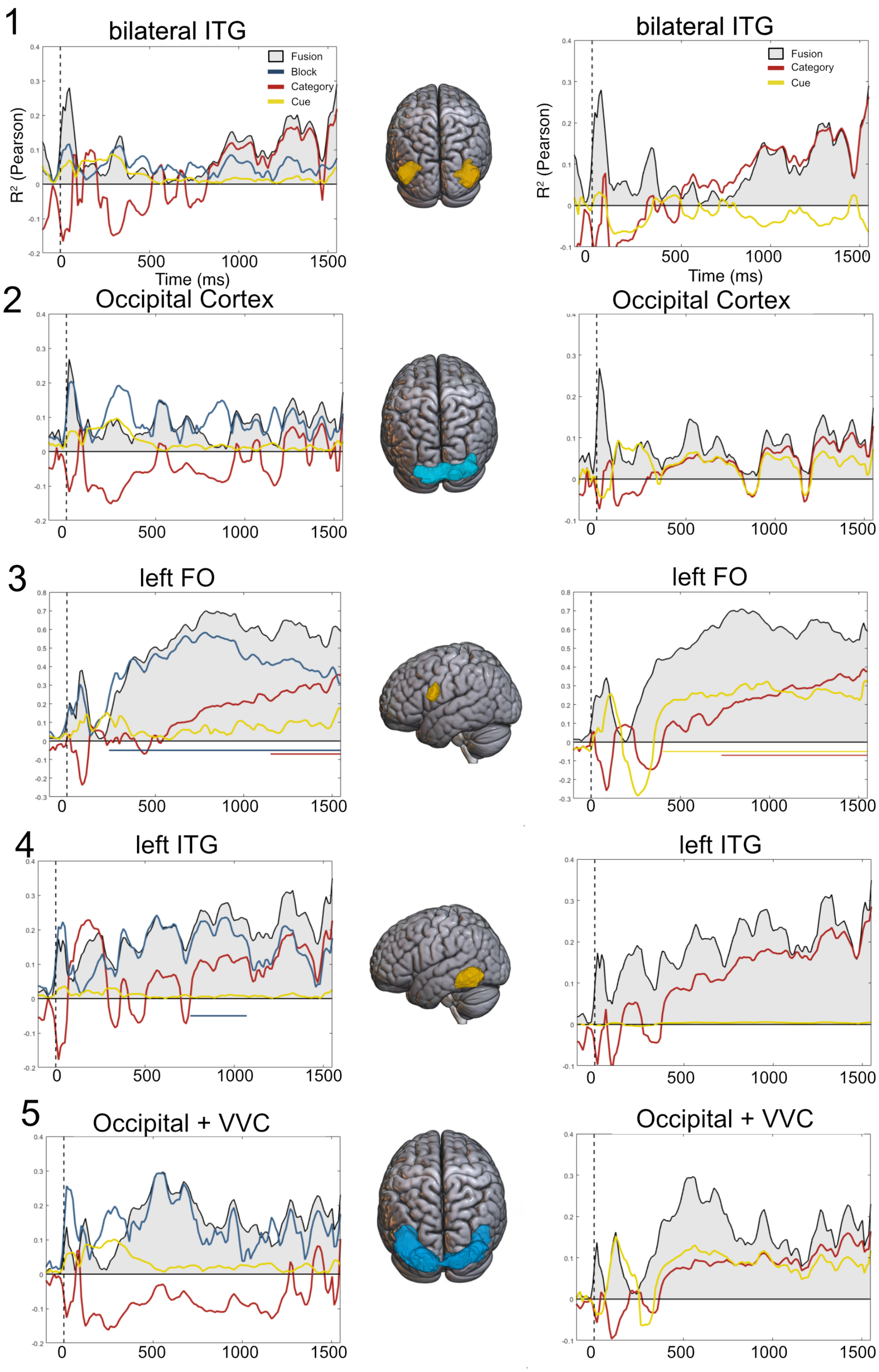


**Supplementary Figure 2.** Fusion values without applying absolute value. ITG = Inferior Temporal Gyrus; FO = Frontal Operculum.

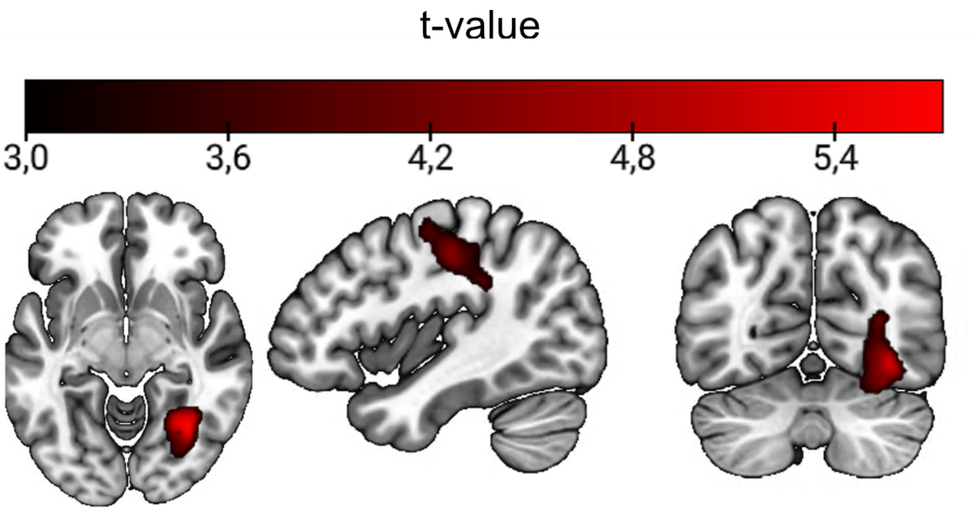


**Supplementary Figure 3.** Cue-category control analyses. Results only shown for the attention condition, since no significant clusters were found in expectation. The classifier was trained to classify between categories using one pair of cues, and then tested in the other.
